# External validation and time-stability analysis of STARE, a blood-free quantification tool for irreversible PET tracers

**DOI:** 10.64898/2026.02.09.704936

**Authors:** Gjertrud L Laurell, Elizabeth A Bartlett, Mike Schmidt, Sergey Anishchenko, Ilia Shkolnik, R Todd Ogden, J John Mann, David Beylin, Jeffrey M Miller, Francesca Zanderigo

## Abstract

**Rationale:** “Gold standard” blood-based quantification of dynamic ^18^F-fluorodeoxyglucose (^18^F-FDG) positron emission tomography (PET) data has limited practical clinical applications due to cost and complexity of data collection and analysis. We previously presented a blood-free quantification alternative, STARE (Source-to-Target Automatic Rotating Estimation), that was validated on ^18^F-FDG data acquired on a ECAT EXACT HR+ scanner. Here, we extend that initial work by externally validating STARE using within-subject data acquired with both a Siemens Biograph mCT scanner and a portable Brain Biosciences CerePET scanner.

**Methods:** Performance was assessed by comparing regional net influx rates (K_i_) estimated using STARE and the standard blood-based Patlak approach. Twenty participants underwent 60-minute ^18^F-FDG scans, on two different days, once in each scanner. The time-stability of both STARE- and Patlak-based K_i_ estimates was evaluated by applying each method to the first 20 (STARE only), 30, 40, and 50 minutes of data.

**Results:** STARE demonstrated high correlation with Patlak K_i_ estimates across both scanner types, particularly in the Biograph mCT (*r* = 0.93), with lower correlation in the CerePET (*r* = 0.71). In the Biograph dataset, STARE provided reliable K_i_ estimates at all evaluated scan durations (20 minutes and above), while in the CerePET dataset, only the 50-minute duration yielded STARE K_i_ estimates that were not significantly different from the full 60 minutes. The Patlak approach provided K_i_ estimates at 40 minutes scan duration and above that did not differ from the 60-min scan results in both datasets.

**Conclusion:** STARE is a viable, noninvasive alternative to traditional blood-based quantification of dynamic ^18^F-FDG PET data, facilitating shorter, blood-free acquisition. This advancement could make dynamic ^18^F-FDG PET imaging more accessible and comfortable for patients, promoting broader clinical adoption.

## INTRODUCTION

Positron emission tomography (PET) allows for quantification of molecular and metabolic processes in the living human brain. Parameters related to underlying biology can be estimated through PET tracer kinetic modeling and graphical approaches (*1*). For PET radiotracers with irreversible pharmacokinetics, metabolic activity can be quantified by estimating the tracer net influx rate (K_i_) into the brain tissue from the vascular compartment (*2*), requiring concurrent measurements of brain tissue radioactivity and plasma tracer concentration to model their relationship. The current “gold standard” for input function estimation relies on measurements of tracer parent radioactivity in arterial plasma, calculated from blood samples collected throughout the scan via arterial cannulation. Although currently considered the closest approximation of the true input to the system, blood-based input functions add complexity, participant burden, and costs to imaging procedures. They are also subject to measurement error.

To address these limitations, efforts have been made to develop, validate, and disseminate less invasive approaches, such as image-derived and population-based input functions, and simultaneous estimation of the input function (*3,4*). These methods reduce reliance on continuous blood sampling, but still require some blood-based “scaling,” making them not completely “blood-free” (*3,4*). Even for tracers like ^18^F-fluorodeoxyglucose (^18^F-FDG), where radiometabolite correction is not necessary for computation of an accurate input function, at least one blood sample is typically required for scaling from whole blood to plasma, or for identification of an input function specific for the subject or scan. Recent developments in image-derived input functions (IDIFs) have aligned with the advent of new, costly, sophisticated, high-resolution scanners with extended field of view. These scanners provide better coverage of major arteries and minimize partial volume effects, better facilitating IDIF approaches (*5*). However, these scanners are unlikely to be widely available in clinical settings in the near future.

The recent introduction of portable, cost-effective PET scanners like CerePET (Brain Bioscience Inc.) offers promising opportunities for both clinical and research applications. Their portability potentially allows for PET scans in remote or rural areas, reducing the need for patients to travel to large hospitals with dedicated PET facilities. Moreover, the lighter weight and flexibility of the scanner enables innovative experimental setups, such as scanning individuals in seated or standing positions, enhancing task-based neuroimaging studies. To ensure true portability of these scanners and enhance applicability of fully quantified PET imaging in general, it is essential to have noninvasive quantification methods that eliminate the need for on-site blood assays.

We previously introduced source-to-target automatic rotating estimation (STARE), the first open-source method to fully quantify PET tracers with irreversible kinetics without relying on any blood data (https://github.com/elizabeth-bartlett/STARE (*6*)). STARE was initially validated against blood-based quantification in ^18^F-FDG PET data acquired using an ECAT EXACT HR+ scanner (*7*–*9*) in a large sample of individuals (N = 69), where STARE-derived K_i_ values were precise and robustly correlated with blood-based K_i_ estimates (*6*).

Here, we applied the same STARE algorithm to analyze new ^18^F-FDG PET data acquired with two different modern scanners. We evaluated STARE’s performance against the blood-based Patlak approach using a recent dataset of 20 healthy subjects imaged with both a standard stationary Biograph mCT (Siemens) and the portable CerePET scanner (*10*). Additionally, we explored the performance of STARE on shorter scan durations, ranging from 20 to 60 minutes, to assess its practicality for clinical application.

## MATERIALS AND METHODS

### Participants

The dataset employed in this study included 20 healthy individuals (10 females; 30.8±10.9 years [range: 18-58]). At the time of imaging, none of the participants were diagnosed with any major psychiatric disorders. Written informed consent was obtained from all participants. The study received approval from the New York State Psychiatric Institute Institutional Review Board and the Joint Radiation Safety Committee at Columbia University. Additional details are available in our previous publication of the data (*10*).

### PET Acquisition and Reconstruction

Each participant underwent two separate 60-minute dynamic ^18^F-FDG PET scans, one with a Biograph mCT (Siemens) scanner and one with a CerePET (Brain Biosciences, Inc.). The scans were scheduled on separate days, with an average inter-scan interval of 57±36 days (range: 1-154 days; CerePET first for 12/20). The tracer was delivered as a 30-second intravenous bolus. During the scans, participants were instructed to rest, keep their eyes closed, and minimize motion. Individualized head molds were used during acquisitions with the Biograph mCT. To mimic realistic use in portable applications, head molds were not used with the CerePET scanner. For both scanners, counts were binned into a total of 22 frames with increasing duration (10×30 s, 5×1 min, 4×5 min, 3×10 min). Maximum-likelihood expectation maximization reconstruction, computed tomography-based attenuation correction, and motion correction were performed as previously described (*10*).

### Blood Data Acquisition

Blood data acquisition for blood-based quantification is described elsewhere (*10*). In brief, arterial blood sampling was completed in n=35 of the total N=40 scans, with collection every 10 seconds for the first 90 seconds and then at 2, 5, 20, 40, and 60 minutes post-injection. For the remaining five scans (2 Biograph, 3 CerePET), venous blood was collected instead at the same time points. Each sample was used to determine the ^18^F-FDG total radioactivity in both plasma and in whole blood.

### Region of Interest Delineation

T1-weighted magnetic resonance (MR) images were acquired for all participants with a 3T GE Signa Premiere scanner. After coregistration, regional time-activity curves (TACs) were extracted using the Desikan-Killany atlas (*11*) available from the FreeSurfer segmentation of each participant’s T1-weighted MR image. TACs from the following eight representative regions were used throughout all analyses: putamen, hippocampus, amygdala, caudal anterior cingulate cortex, lateral occipital cortex, middle temporal cortex, rostral middle frontal cortex, and superior parietal cortex, as done with previous analysis of the same data (*10*).

### Blood-Based Quantification

Arterial input functions were derived from measures of tracer radioactivity in arterial plasma. The data were linearly interpolated between the start of the scan and the peak plasma activity, while a tri-exponential decay function was fitted from the peak plasma activity to the end of the scan. For scans with venous samples, input functions were generated by scaling a template input function – obtained as the average of all the available arterial input functions in the sample, once expressed in standard uptake value units – to match the radioactivity in venous plasma at 40 minutes post-injection, where ^18^F-FDG levels in venous and arterial blood equilibrate (*12*).

Using the input functions, regional K_i_ values were estimated via the Patlak approach (*13*) with a fixed start of the linear phase (t*) of 17.5 min, across all scans and ROIs. Vascular correction was performed with a fixed fractional blood volume (v_B_) of 5% across all ROIs and scans, using the measured total ^18^F-FDG radioactivity in whole blood modeled using the same approach as the plasma data.

### Blood-Free Quantification with STARE

STARE estimates regional K_i_ values from individual PET data without blood radioactivity measurements (*6*). The algorithm models tracer dynamics in multiple target regions based on tracer concentration in another (“source”) region, thus eliminating the need for an arterial input function, and simultaneously estimates system parameters for target and source regions. Each ROI is considered as a source in turn, with K_i_ estimates for each target region averaged from all source contributions. To ensure identifiability, STARE employs an anchoring term and free parameter bounds, both derived automatically from the PET data. For this purpose, a two-step k-means clustering approach is used to identify voxels in the field of view likely representing vasculature, which subsequently undergo partial volume correction (PVC). The vascular signal is then bootstrapped, simulating curves within one standard deviation above and below the vascular cluster TAC. A two-tissue compartment irreversible model is then fitted using these bootstrapped curves as input function proxies, generating distributions of free parameters. From these, each parameter’s bounds are set at the distributions’ full width at half maximum (FWHM). The midpoint of the K_i_ bounds serves as an anchoring term in the STARE cost function.

We applied the publicly available STARE algorithm (https://github.com/elizabeth-bartlett/STARE), implemented in Matlab (version 9.1), without alterations from the version used in our previous study (*6*). All eight ROIs listed above were included in the analysis. Results from all source contributions were averaged with equal weighting across ROIs, to provide the final K_i_ estimates. For the main analyses, no vascular correction was applied (v_B_set to 0% across ROIs). However, for the Biograph mCT data with full scan duration, we conducted a comparative analysis using STARE with vascular correction (v_B_= 5%), employing the PVCed vascular cluster TACs as a proxy for radioactivity in whole blood (*6*).

The required user input for the FWHM of the point spread function (PSF) in the PVC step is estimated according to 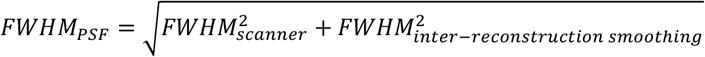 Here, *FWHM _scanner_* is the FWHM scanner spatial resolution at the center of the field of view, and *FWHM _inter-reconstruction smoothing_* is the FWHM of the spatial smoothing filter applied in image reconstruction.

For the Biograph mCT scans, the bottom six slices of all images were removed prior to vascular clustering to reduce effects of noise, as done previously (*6*). In the PVC step, a PSF with 5.9 mm FWHM was used.

For the CerePET scans, clipping a fixed number of slices was not feasible, due to the shorter axial field of view and large variations in brain positioning across participants. Instead, the vascular clustering step was initially run on full images. The position of the clusters and the shape of the resulting centroid were then visually inspected, and 15 of the 20 scans deemed to produce adequate clusters. For the remaining five, the analysis was repeated with images cropped at four slices below the bottom of the cerebellum (between 5 and 21 slices clipped) to obtain reasonable vascular clusters. In the PVC step, a PSF with 4.1 mm FWHM was used.

### Time-Stability Analyses

To assess the time-stability of K_i_ estimates, we applied Patlak and STARE to the first 50, 40, and 30 minutes of the data post-injection, by sequentially removing the later scan frames. Reducing the scan duration below 30 minutes was not feasible with Patlak, due to limited data in the linear phase, but STARE was additionally tested using only the first 20 minutes of each scan.

For these analyses, scans without arterial samples were excluded, yielding 18 Biograph mCT scans and 17 CerePET scans. For the Patlak approach, new arterial input functions were generated by fitting the blood data collected within each abbreviated duration. STARE was applied consistently across scan durations, including keeping clipping positions determined from the full scan data as outlined above.

### Statistical Analysis

Percent differences in regional K_i_ estimates were calculated as 100 · (*K_i, STARE_* − *K_i, Patlak_*)/ *average*(*K_i, STARE_* − *K_i, Patlak_*) for between-method differences and as 100 · (*K_i, CerePET_* − *K_i, Biograph_*)/*average*(*K_i, CerePET_* − *K_i, Biograph_*) for between-scanner differences. We report mean and standard deviations of the percentage differences across all scans and regions. To assess within-participant and within-region correspondence of K_i_ estimates between analytical methods (Patlak, STARE) and scanners (Biograph, CerePET), simple linear regression was employed. We report slopes, intercepts and Pearson’s correlation coefficients (*r*). Linear mixed effects models were applied to log-transformed K_i_ estimates (to satisfy the requirements of the model) to assess method (STARE, Patlak) and scanner (CerePET, Biograph) differences. ROI, as well as method or scanner respectively, were included as fixed effects and participant as a random effect.

For visualization of time-stability, we computed the percentage bias of the K_i_ estimates at each scan duration relative to K_i_ values estimated from 60 minutes of data as 100 · (*K*_*i, x* 60 *mins*_ − *K_i,_* _60 *mins*_)/*K_i,_* _60 *mins*_. We report median bias and median absolute deviation (MAD) of the bias across all regions and participants. Linear mixed effects models were applied to log-transformed K_i_ estimates to assess within-method differences between scan durations, with ROI included as a fixed effect, and participant as a random effect.

Linear mixed effects models were calculated with R (version 4.4.1), and all other analyses were run with Matlab (version 9.1).

## RESULTS

### STARE vs. Patlak

Across all ROIs and scans, the mean difference between blood-free STARE and blood-based Patlak K_i_ estimates was -1.11%±12.11% for Biograph mCT and 0.83%±22.74% for CerePET, though between-method differences were not statistically significant in either dataset (p = 0.684 and p = 0.767, respectively). STARE K_i_ estimates exhibited strong correlations with Patlak K_i_ estimates in both scanners (Figure 1). Within-participant, across-regions regression slopes ranged from 0.60 to 1.12 (*r*: 0.99-1.00) in Biograph, and from 0.68 to 1.57 (*r*: 0.78-1.00) in CerePET, with individual regression parameters presented in Figure 1. Within-region, across-participants regression slopes ranged from 1.01 to 1.08 (*r*: 0.82-0.93) in Biograph, and from 0.36 to 0.77 (*r*: 0.20-0.66) in CerePET, with individual parameters presented in Tables 1 and 2. Including vascular correction in the STARE algorithm did not improve correlation with Patlak K_i_ values (Supplementary Figure 1).

**Table 1:** Within-region, across-participant comparison of STARE and Patlak K_i_ values in the Biograph mCT data.

| | Regression slope | $r$ |
| --- | --- | --- |
| Amygdala | 1.03 | 0.85 |
| Caudal anterior cingulate cortex | 1.08 | 0.93 |
| Hippocampus | 1.01 | 0.83 |
| Lateral occipital cortex | 1.08 | 0.82 |
| Middle temporal cortex | 1.04 | 0.89 |
| Putamen | 1.04 | 0.89 |
| Rostral middle frontal cortex | 1.02 | 0.91 |
| Superior parietal cortex | 1.08 | 0.85 |

**Table 2:** Within-region, across-participant comparison of STARE and Patlak K_i_ values in the CerePET data

| | Regression slope | $r$ |
| --- | --- | --- |
| Amygdala | 0.77 | 0.41 |
| Caudal anterior cingulate cortex | 1.08 | 0.66 |
| Hippocampus | 0.68 | 0.38 |
| Lateral occipital cortex | 0.68 | 0.39 |
| Middle temporal cortex | 0.84 | 0.46 |
| Putamen | 0.85 | 0.47 |
| Rostral middle frontal cortex | 0.77 | 0.39 |
| Superior parietal cortex | 0.36 | 0.20 |

**Figure 1.**
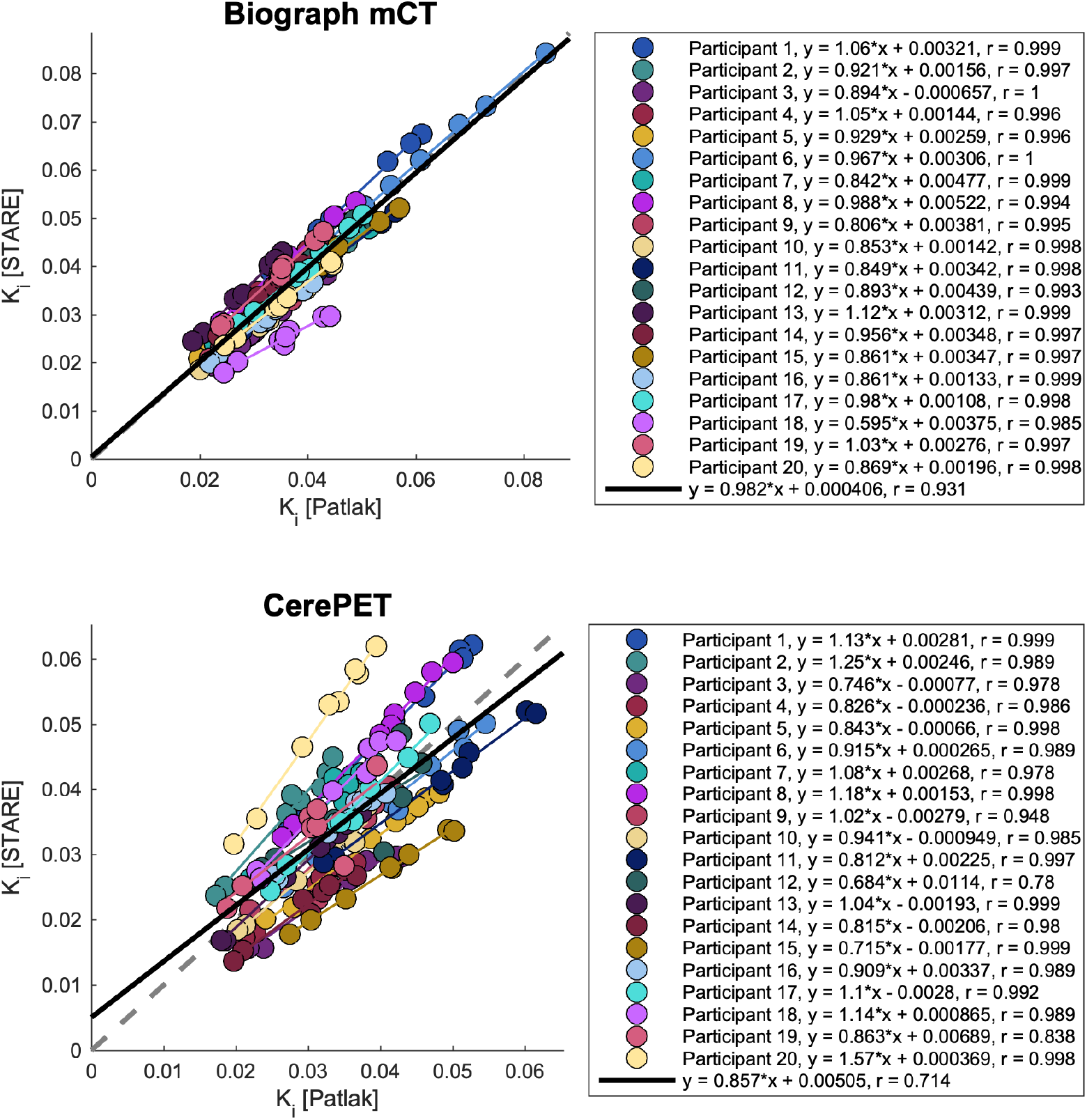
STARE (blood-free) vs. Patlak (blood-based) K_i_ for Biograph mCT (top) and CerePET (bottom). Individual regression lines are plotted in each participant’s unique marker color. Within-scan, across-regions regression parameters are reported in the legend. The grey dotted line represents the line of identity.

### CerePET vs. Biograph mCT

K_i_ values were overall lower in the CerePET scans relative to the Biograph mCT scans, both when using Patlak, as reported previously (*10*), and STARE (-3.39%±26.18%), though main effect of scanner was not statistically significant with either method. None of the statistically significant region-by-scanner interactions previously found with Patlak (*10*) were observed with STARE K_i_ values. The correlation between corresponding CerePET and Biograph K_i_ values was lower for STARE (*r* = 0.61: Figure 2), than we previously reported for Patlak (*10*) (*r* = 0.65). For STARE, within-participant, across-regions regression slopes ranged from 0.43 to 1.72 (*r*: 0.61-0.99), with individual regression parameters reported in Figure 2. Within-region, across-participants regression slopes ranged from 0.29 to 0.53 (*r*: 0.29-0.52), with individual parameters presented in Table 3. Equivalent figures and tables for Patlak K_i_ estimates are previously published (*10*). As previously found with Patlak, between-scanner agreement was improved when Participant 6, who had unusually high K_i_ estimates in the Biograph scan, was excluded from the analyses (*r* = 0.65; Supplementary Figure 2).

**Table 3:** Within-region, across-participant comparison of CerePET and Biograph mCT STARE K_i_ values.

| | Regression slope | $r$ |
| --- | --- | --- |
| Amygdala | 0.31 | 0.29 |
| Caudal anterior cingulate cortex | 0.47 | 0.52 |
| Hippocampus | 0.53 | 0.48 |
| Lateral occipital cortex | 0.39 | 0.30 |
| Middle temporal cortex | 0.39 | 0.37 |
| Putamen | 0.49 | 0.48 |
| Rostral middle frontal cortex | 0.29 | 0.34 |
| Superior parietal cortex | 0.35 | 0.34 |

**Figure 2.**
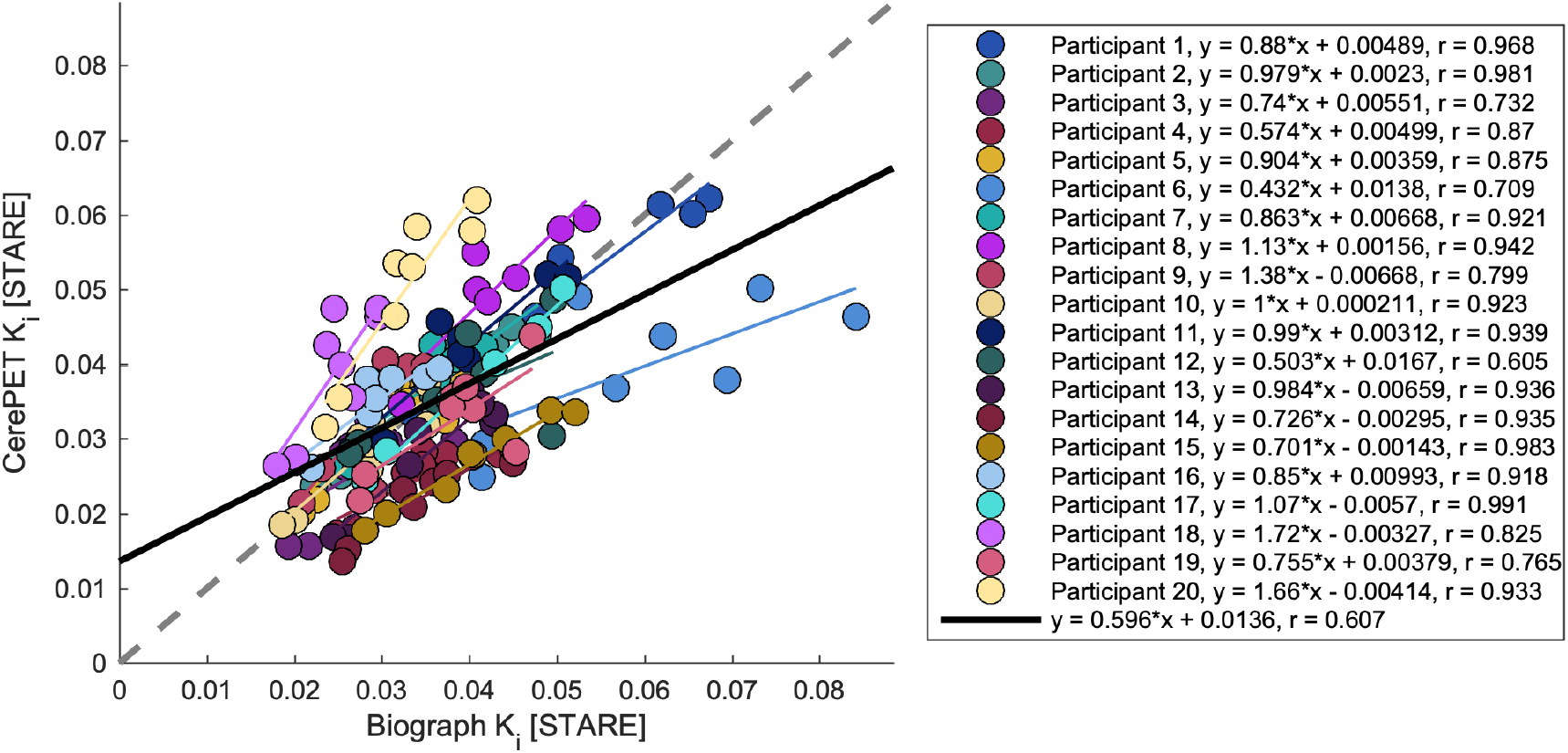
CerePET K_i_ values vs Biograph mCT K_i_ values, both estimated with STARE. Within-participant, across-regions regression parameters are reported in the legend. The grey dotted line represents the line of identity.

### Time-stability

For STARE, the time-stability was markedly superior in the Biograph mCT dataset compared to the CerePET (Figure 3, Table 4). For Patlak, there was less difference in performance between scanners, although the sign of the median bias at 30-minute scan duration was positive (15.6%±8.2%, p = 0.01) in the Biograph mCT dataset and negative (-6.9%±17.9%, p = 0.24) in the CerePET dataset (Figure 3, Table 5). For STARE on the Biograph data, none of the scan durations produced K_i_ estimates that were significantly different from those estimated from the 60-minute scan. In the CerePET, however, STARE results were significantly different at 40 minutes and below. For Patlak on the Biograph data, the K_i_ estimates were significantly different at 30 minutes. For Patlak on the CerePET data, although none of the considered durations produced K_i_ estimates significantly different from those calculated based on the 60-minute scan duration, the bias changed directionality between the 40-and 30-minute durations, and there was a relatively high variability in the bias at 30 minutes (Table 5). For STARE, especially in the Biograph mCT data, the strong agreement with Patlak K_i_ estimated from the full scan duration remained across all abbreviated scan durations (Figure 4).

**Table 4:** Comparison of STARE K_i_ values estimated using shorter scan durations to those derived from 60 minutes of data. p-values indicate results of linear mixed effects models testing for effect of scan duration.

| Duration | Biograph |  | CerePET |  |
| --- | --- | --- | --- | --- |
|  | Bias | p-value | Bias | p-value |
|  | [median±MAD] |  | [median±MAD] |  |
| 50 min | 1.81%±5.34% | 0.991 | 9.67%±11.27% | 0.853 |
| 40 min | 0.99%±5.68% | 0.978 | 14.55%±12.62% | 0.007 |
| 30 min | 2.05%±7.81% | 0.886 | 28.43%±19.98% | <0.0001 |
| 20 min | 2.79%±6.37% | 0.481 | 41.64%±28.90% | <0.0001 |

MAD = median absolute deviation

**Table 5:** Comparison of Patlak K_i_ values estimated using shorter scan durations to those derived from 60 minutes of data. p-values indicate results of linear mixed effects models testing for effect of scan duration.

| Duration | Biograph |  | CerePET |  |
| --- | --- | --- | --- | --- |
|  | Bias |  | Bias |  |
|  | [median±MAD] | p-value | [median±MAD] | p-value |
| 50 min | 3.94%±2.06% | 0.608 | 2.31%±2.49% | 0.995 |
| 40 min | 7.45%±2.15% | 0.134 | 8.57%±4.26% | 0.310 |
| 30 min | 15.61%±8.23% | 0.014 | -6.86%±17.88% | 0.244 |

MAD = median absolute deviation

**Figure 3.**
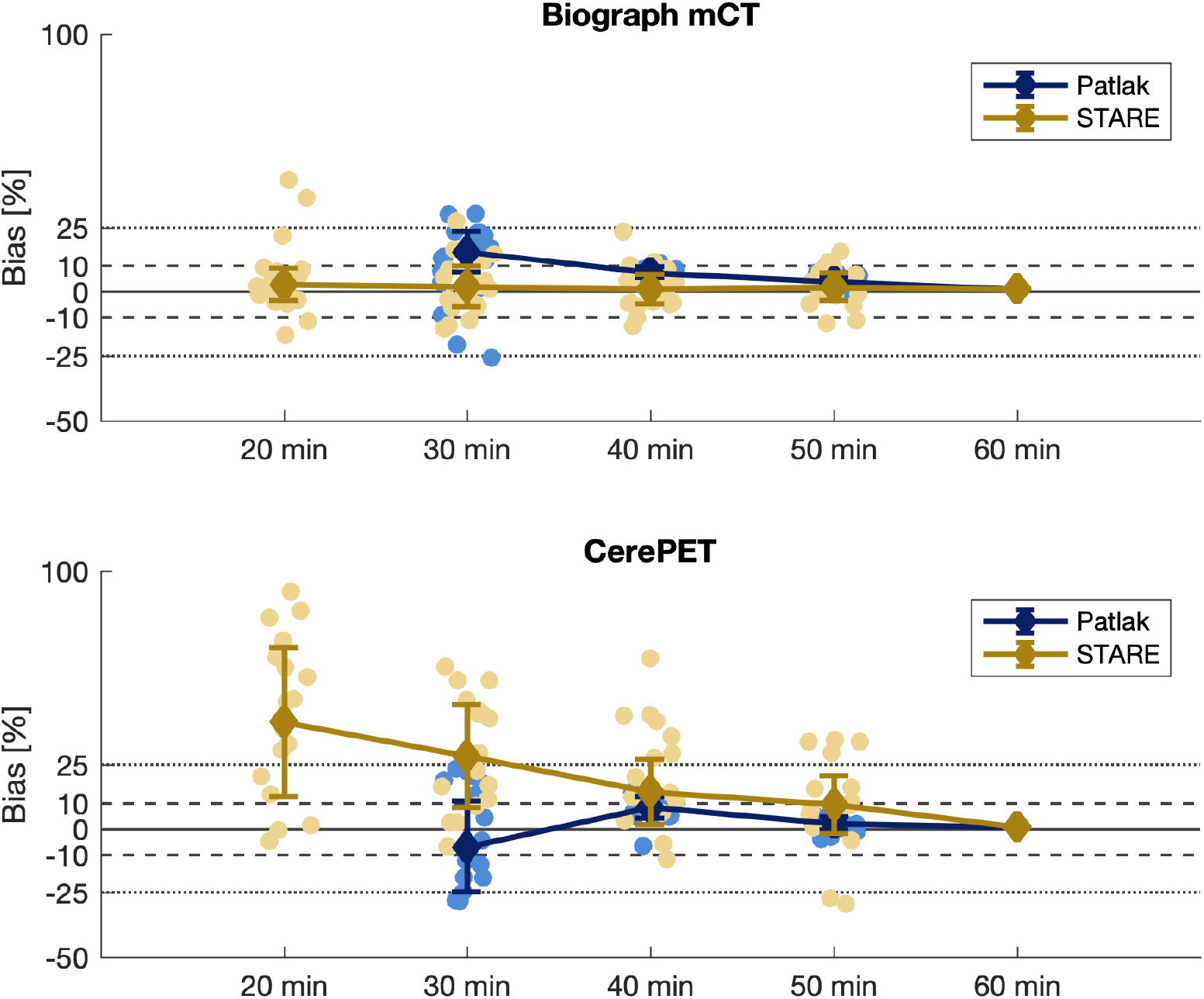
Median bias in K_i_ estimates compared to 60-minute scan duration for STARE and Patlak in the Biograph mCT (top) and CerePET (bottom) datasets. Error bars denote the median absolute deviation, and circular markers denote the across-regions median bias for each participant.

**Figure 4.**
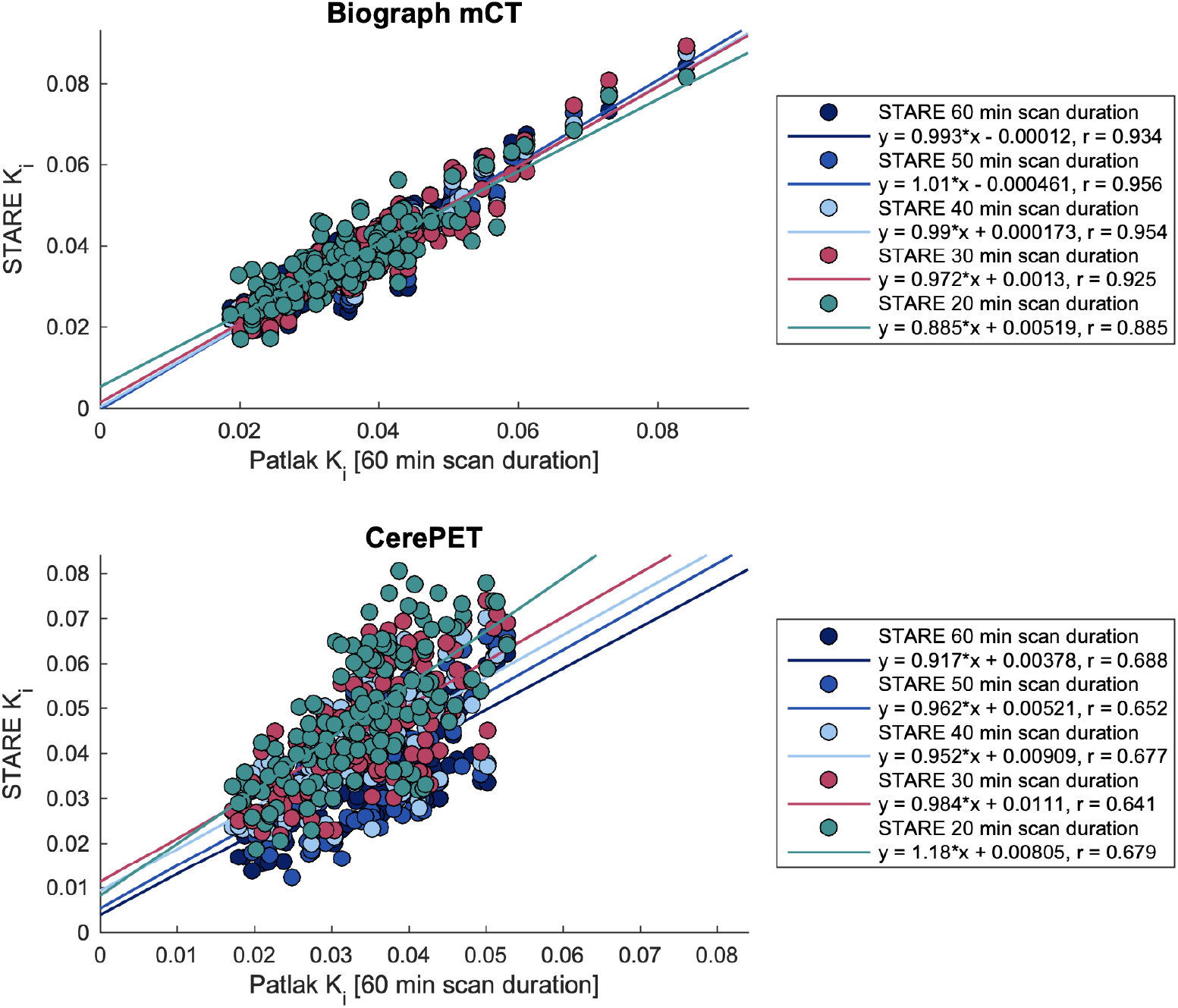
STARE K_i_ estimates from different scan durations plotted against corresponding Patlak K_i_ values estimated from the full 60 minutes of data. All ROIs and all participants with arterial blood (n=18 for Biograph; n=17 for CerePET) are included.

## DISCUSSION

This study extends the application of STARE, initially only validated with ^18^F-FDG data acquired from an ECAT EXACT HR+ scanner, to a recent dataset of ^18^F-FDG PET scans acquired within-subject using both a clinical Biograph mCT scanner and the portable CerePET scanner. Despite differences in scanner technology, population age, disease state, and image reconstruction methods between the previous study and the current one, we again observe strong correlations between STARE and blood-based estimates of K_i_. STARE showed strong correlation with blood-based Patlak K_i_ estimates, particularly in the Biograph mCT data, where the correlation coefficient across regions and participants (0.93) exceeded the previously reported value (0.83) when comparing STARE to two-tissue irreversible compartment model-based K_i_ estimates from ECAT EXACT HR+ ^18^F-FDG PET scans (*6*). These findings across two distinct independent datasets strengthen our confidence in recommending STARE as a viable alternative to traditional blood-based quantification of ^18^F-FDG PET in clinical scanners.

Our findings demonstrate that STARE can accurately estimate K_i_ without the need for blood sampling even at shorter scan durations. This is crucial for clinical implementation, as shorter scan times enhance patient comfort and feasibility, and reduce costs. Full quantification is rarely employed clinically, as it typically requires both continuous blood sampling and relatively long scan durations. Instead, semi-quantitative outcome measures like the standardized uptake value or standardized uptake value ratios are frequently used. While these outcomes significantly reduce the cost and complexity of PET acquisition and analyses, they are associated with challenges such as lower sensitivity, lack of specificity and possible confounding of outcomes by global changes or use of improper reference regions (*14*).

Although STARE performed well across both scanners, the lower correlation and poorer time-stability observed in the CerePET suggests room for improvement for both the CerePET hardware and the STARE algorithm. The lower correlation in the CerePET data may stem from its operational mechanics. Specifically, the detector rings, which do not encompass the entire 22.5-cm axial field of view, move during scanning to cover the brain. Thus, activity from different parts of the brain is detected at different times within each frame, affecting the uniformity of tracer delivery detection across the brain and generating overall noisier TACs. The next-generation CerePET scanner has full brain coverage with static rings, allowing for more accurate characterization of tracer delivery, which is expected to lead to improved STARE performance. As an alternative to hardware updates, a more sophisticated reconstruction approach could be implemented, accounting for the movement of the detectors during acquisition. It is also possible that the lack of head molds during acquisition with CerePET could also contribute to inter- and intra-frame head motion and blurring. As previously reported, there was on average more inter-frame participant movement detected by the motion correction algorithm in the CerePET scans compared with scans acquired in the Biograph mCT, although the difference was not statistically significant (*10*). To enhance the robustness of STARE, particularly in the vascular cluster extraction step, we are currently developing improvements to the algorithm, such as integrating cluster sparsity measures and centroid signal variance into the cluster selection process. These planned enhancements, alongside anticipated hardware upgrades to the CerePET, are expected to enhance STARE’s performance on CerePET data. Successful integration of STARE with CerePET data at shorter scan durations would increase the application potential of this type of scanner.

In the Biograph mCT dataset, STARE showed excellent time-stability, with K_i_ values estimated from 20 minutes of data not significantly different from 60 minutes (Table 3), and still strongly correlated with Patlak K_i_ values from the full scan duration (Figure 4). For Patlak, the shortest scan duration in the Biograph that was not significantly different from the full scan duration was 40 minutes. The data binning was based on a 60-minute protocol, and it is possible that a 30-minute acquisition with shorter frames between 17.5 and 30 minutes would have led to better Patlak performance than observed here (15.61±8.23% positive bias). Positive K_i_ bias with shorter scan durations, for both the two-tissue irreversible model and Patlak approach, have previously been reported for tumor lesions outside of the brain (*15,16*). Though we observe a larger bias than those previously reported, it has also been shown that abbreviated scan protocols with Patlak might be more challenging in brain than in the periphery (*17*).

Here, we have demonstrated STARE’s utility for application to ^18^F-FDG data acquired on traditional clinical scanners with abbreviated scan protocols as short as 20 minutes, and for ^18^F-FDG data acquired on the current generation of CerePET scanners with 60 minutes scan duration. For broader application of STARE, such as application to other tracers with irreversible kinetics or to CerePET data with shorter acquisition times, further optimization and validation of the algorithm is required. This includes determining the number of ROIs to include and the weighting of the anchoring term. Future work will include extension to voxel-level analyses and addressing metabolite correction for irreversible tracers with radioactive metabolites (e.g., ^11^C-arachidonic acid tracers, ^18^F-FMISO, ^11^C-choline, and ^11^C-acetate). A Python implementation of STARE is currently under validation.

## CONCLUSION

Our findings underscore the transformative potential of STARE in simplifying and shortening ^18^F-FDG PET imaging protocols. STARE offers a versatile and efficient solution for obtaining reliable K_i_ estimates in clinical settings. As we continue to refine and validate STARE across scanners and settings in further external validations, its integration with technologies like CerePET holds the promise of making fully quantified ^18^F-FDG PET imaging truly portable.

## DISCLOSURE

The National Institute of Biomedical Imaging and Bioengineering funded this study (R01EB026481; principal investigator, Francesca Zanderigo). J. John Mann receives royalties from the Research Foundation for Mental Hygiene for the commercial use of the Columbia Suicide Severity Rating Scale. Ilia Shkolnik (senior software developer), Sergey Anishchenko (computational scientist), and David Beylin (chief executive officer) have financial interests as employees and shareholders at Brain Biosciences. No other potential conflict of interest relevant to this article was reported.

## Notes

### Competing Interest Statement

Ilia Shkolnik, Sergey Anishchenko, and David Beylin have financial interests as employees and shareholders at Brain Biosciences. No other potential competing interests relevant to this article was reported.

