## Supplemental Figures for "External validation and time-stability analysis of STARE, a blood-free quantification tool for irreversible PET tracers"

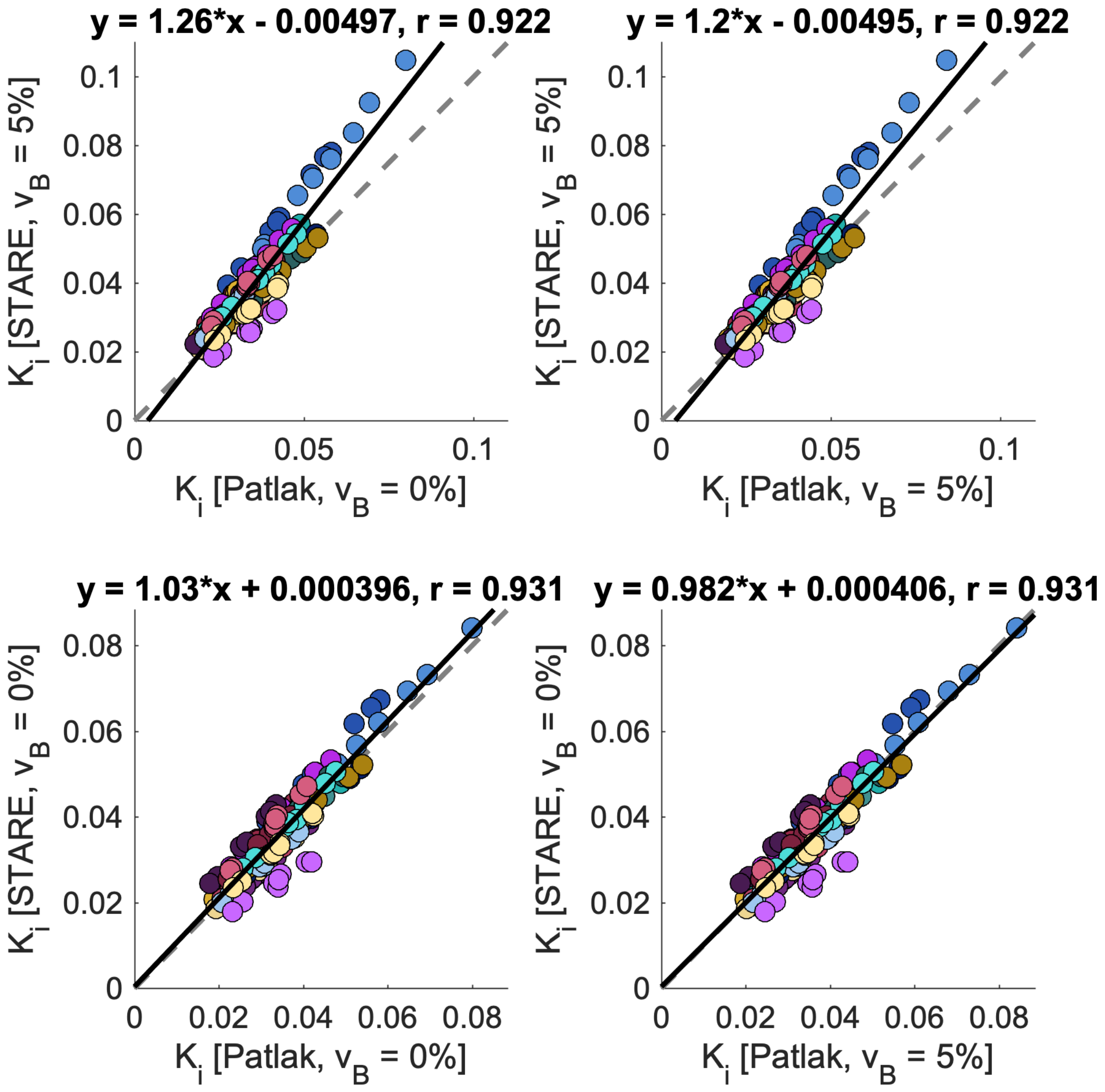


Supplemental Figure 1: Blood-free Ki estimates from STARE plotted against blood- based Ki estimates from Patlak for the Biograph mCT scans, for both methods, both with and without vascular correction.

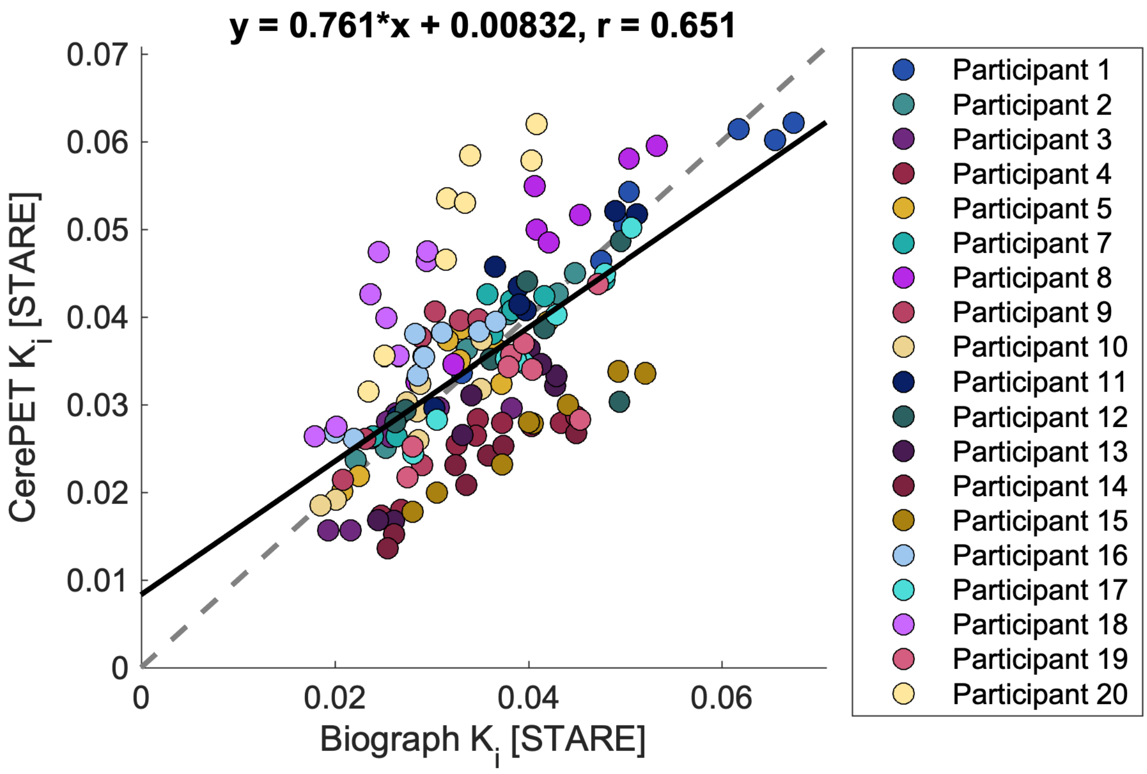


Supplemental Figure 2: STARE Ki estimates from CerePET scans plotted against STARE Ki estimates from the corresponding Biograph mCT scans, excluding participant 6.
